# Peripubertal antagonism of corticotropin-releasing factor receptor 1 results in sustained changes in behavioral plasticity and the transcriptomic profile of the amygdala

**DOI:** 10.1101/2024.08.14.607957

**Authors:** Julia Martz, Micah A. Shelton, Tristen J. Langen, Sakhi Srinivasan, Marianne L. Seney, Amanda C. Kentner

**Author notes:** Corresponding author: Amanda C. Kentner, Office.

## Abstract

Peripuberty is a significant period of neurodevelopment with long-lasting effects on the brain and behavior. Blocking type 1 corticotropin-releasing factor receptors (CRFR1) in neonatal and peripubertal rats attenuates detrimental effects of early-life stress on neural plasticity, behavior, and stress hormone action, long after exposure to the drug has ended. CRFR1 antagonism can also impact neural and behavioral development in the absence of stressful stimuli, suggesting sustained alterations under baseline conditions. To investigate this further, we administered the CRFR1 antagonist (CRFR1a) R121919 to young adolescent male and female rats across 4 days. Following each treatment, rats were tested for locomotion, social behavior, mechanical allodynia, or prepulse inhibition (PPI). Acute CRFR1 blockade immediately reduced PPI in peripubertal males, but not females. In adulthood, each assay was repeated without CRFR1a exposure to test for persistent effects of the adolescent treatment. Males continued to experience deficits in PPI while females displayed altered locomotion, PPI, and social behavior. The amygdala was collected to measure long-term effects on gene expression. In the adult amygdala, peripubertal CRFR1a induced alterations in pathways related to neural plasticity and stress in males. In females, pathways related to central nervous system myelination, cell junction organization, and glutamatergic regulation of synaptic transmission were affected. Understanding how acute exposure to neuropharmacological agents can have sustained impacts on brain and behavior, in the absence of further exposures, has important clinical implications for developing adolescents.

## Introduction

Corticotropin-releasing factor (CRF) is a major regulator of the stress response through its actions on the hypothalamic-pituitary-adrenal (HPA) axis and its associated receptors (CRFR) type 1 and 2 (Binder and Nemeroff, 2010). CRF works in brain regions such as the amygdala and hippocampus to alter neural plasticity, learning and memory, and future responses to stressors (Vandael et al., 2021). Stress exposure during sensitive periods of early development can have long-lasting effects, including the upregulation of associated CRF receptors in the brain (Veenit et al., 2014; Núñez Estevez et al., 2020; Zhao et al., 2021). Other effects include an increased risk of developing cognitive deficits and psychopathologies, such as anxiety disorder and schizophrenia in humans (Carr et al., 2013; Smith and Pollak, 2020) and associated symptoms in other animals (Fenoglio et al., 2005; Wang et al., 2012; Toth et al., 2015; Veenit et al., 2014; Hu et al., 2020; Núñez Estevez et al., 2020; Zhao et al., 2021). Treatment with CRFR1 antagonists (CRFR1a) at sensitive periods, such as during neonatal development or in peripuberty, can reduce neural and behavioral impairments in animal models of early life stress (Ivy et al., 2010; Short et al., 2020; Veenit et al., 2014). In these studies, a week-long exposure to the drug attenuated hippocampal deficits in long-term potentiation (LTP), prevented hippocampal increases in CRF, and mitigated associated changes in social, affective, and cognitive functioning in male rodents. Notably, these rescue effects were observed months after treatment exposure. Peripubertal CRFR1 blockade alone, in the absence of a negative experience also affected the stress response in later life (Veenit et al., 2014). This highlights that CRFR1 deficiency, and not only upregulation, can shape behavioral and cognitive functioning. Together, these data are suggestive of sustained changes in neural- and behavioral plasticity following alterations to CRFR1 functioning during discrete developmental periods such as puberty.

The short- and long-term behavioral changes brought on by exposure to new experiences and environments are the result of neuroplasticity, which occurs when the brain changes how it functions in response to these novel experiences. Stress-related brain systems are particularly sensitive to these changes in synaptic function and plasticity (Chen et al., 2012; Dos-Santos et al., 2023). For example, CRF acting in the amygdala has been shown to enhance excitability of CRFR1 neurons and trigger changes in neural signaling, leading to long-lasting changes in sensorimotor gating, anxiety-like behavior, and fear discrimination (Rajbhandari et al., 2015; Rainnie et al., 2004; Sanford et al., 2017). Neural and behavioral alterations are facilitated by expression changes in genes related to plasticity, such as those encoding for Eph receptor A4 and adenosine receptor A2a ([*Epha4]* Deininger et al., 2008; [*Adora2a]* Simões et al., 2016), as well as genes associated with neurodevelopmental disorders (glial fibrillary acidic protein [*gfap*] and teashirt zinc finger homeobox *[Tshz3]*; Herrero et al., 2020; Caubit et al., 2016) and stress (CRF receptors *Crhr1 and Crhr2*). Furthermore, CRF has been shown to directly regulate synaptic function and subsequent behavioral outcomes through activation of cannabinoid receptors (Jamieson et al., 2022; Ruat et al., 2021). These stress-sensitive pathways are particularly malleable during sensitive periods of development such as early adolescence (Filetti et al., 2023). Therefore, while previous work focused on social, cognitive, and affective measures of behavioral plasticity (Ivy et al., 2010; Short et al., 2020; Veenit et al., 2014), we extended our study to include measures of mechanical allodynia, locomotor behavior, and prepulse-inhibition (PPI) which are also influenced by CRF functioning in the amygdala (Carty et al., 2010; Chang et al., 2022; Cui et al., 2004; Liang et al., 1992; Sutherland et al., 2010; Rajbhandari et al., 2015; Bijlsma et al., 2011; Tian et al., 2024).

In the absence of stressors, studies in adult rodents have found that neither acute nor chronic blockade of CRFR1 affected circulating levels of corticosterone or adrenocorticotropic hormone ([ACTH] Keck et al., 2001; Gutman et al., 2011). However, while acute CRFR1a administration had no effect on sensorimotor gating as measured by the extent of pre-pulse inhibition ([PPI)] Sutherland and Conti, 2011), chronic infusions of CRFR1a have been shown to reduce anxiety-like behaviors in defensive withdrawal tasks in control animals that were not exposed to stressful stimuli (Arborelius et al., 2000; Gutman et al., 2011). Gutman and colleagues (2011) also observed that the density of CRF mRNA in the central nucleus of the amygdala (CeA) decreased after chronic blockade of CRFR1, which was likely tied to reductions in anxiety-like behavior (McCall et al., 2015). Taken together, these data suggest that CRFR1a may modulate some behaviors under baseline conditions when a stressful stimulus is not present. However, these studies only evaluated adult male rats and did not consider the influences of various developmental periods. Stress, or lack thereof, has been shown to influence the brain more prominently during early life and during peripuberty, compared to adulthood (Kirby et al., 2013; Lee and Jung, 2024). More research is needed to understand how acute and chronic blockade of CRFR1 at discrete time points across development might affect behavioral function in both male and female animals.

Although CRFR1a prevents adverse effects associated with a stressful experience (Ivy et al., 2010; Short et al., 2020; Veenit et al., 2014), it is also important to understand how altered functioning of CRF receptors may affect neural plasticity and behavior in the absence of explicitly stressful stimuli. Indeed, individuals can receive neuropharmacological interventions during acute, chronic, or even under low or “regular” stress conditions. We aim to identify underlying mechanisms of how exposure to psychoactive agents at defined developmental periods can continue to influence brain function and behavior without further drug exposure. This is relevant for CRFR1 antagonists where there has been a sustained interest for their clinical potential amongst the scientific community (Alizamini et al., 2023; Künzel et al., 2003; Lv et al., 2022; Schreiber et al., 2018; Zorrilla & Koob, 2010). To this end, we acutely administered a CRFR1a, without an accompanying stressor, to male and female rats across 4 days in early adolescence. One hour after each drug treatment, rats were tested for locomotor behavior, social preference, social discrimination, mechanical allodynia, or PPI. These behavioral tests were repeated in adulthood, but without CRFR1a treatment, and amygdala samples were collected to assess whether blockade of CRFR1 during early adolescence imparts long-term changes in genes related to neural plasticity and stress responses.

## Experimental Procedures

### Animals and Experimental Overview

Male and female Sprague Dawley rats were purchased from Charles River Laboratories (Males: Wilmington, MA; Females: Raleigh, NC) and housed with *ad libitum* access to food and water on a light/dark cycle (0700-1900 light) at 20°C. After habituating to the facility, animals were bred (1 male to 2 females) and pregnancy confirmed by the visualization of spermatozoa in vaginal samples, a sustained diestrus phase, and constant body weight gain. In line with the 3Rs of animal research’s principle of *Reduction* (NC3Rs, n.d), the experimental subjects from this study (run across multiple cohorts) were siblings retained at weaning from a previous maternal immune activation study (Martz et al., 2024) whose mothers were control dams treated with saline on gestational day 15. All animal procedures were approved by the Massachusetts College of Pharmacy and Health Sciences (MCPHS) Institutional Animal Care and Use Committee and were carried out in compliance with the Association for Assessment and Accreditation of Laboratory Animal Care. The experimental procedures and timeline are presented in **Fig. 1A**.

**Figure 1.**
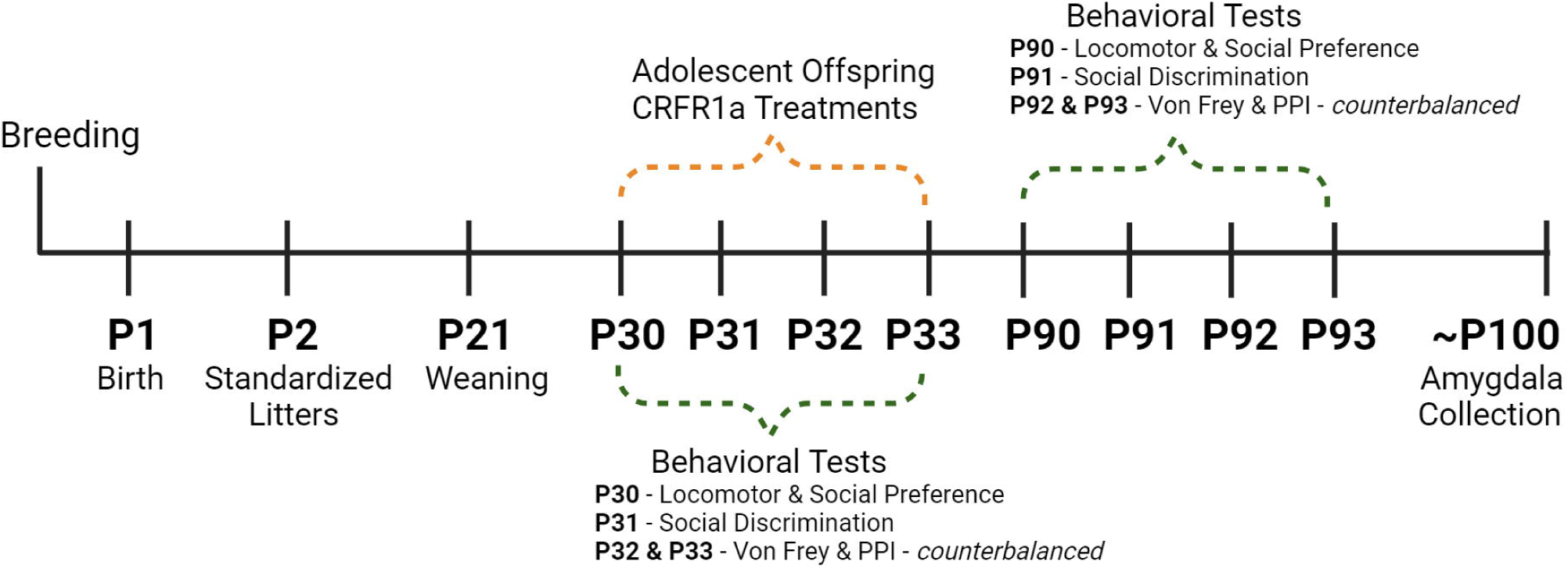
Experimental Timeline.

Day of birth was designated as postnatal day (P)1 and offspring were culled to 10 pups per litter (5 males and 5 females, wherever possible), staying with their mothers until weaning on P21. At that point, offspring were placed into clean cages in same-sex pairs where they were maintained throughout the rest of the study. Offspring were treated with a daily subcutaneous injection of either the selective CRFR1a R121919 (CRFR1a; 10mg/kg, MCE MedChemExpress, Monmouth, NJ) or pyrogen free saline (vehicle control), 1-hour prior to behavioral testing, across a 4-day period. Specifically, on P30, animals received either CRFR1a or vehicle exposure and were evaluated on their locomotor activity followed by a social preference test. On P31 animals were evaluated on their social discrimination ability. Prepulse inhibition (PPI) and von Frey tests were counterbalanced between P32 and P33. We selected a dose of 10 mg/kg because others have reported behavioral effects with repeated administration of R121919 at this dose, administration route, and behavioral testing timeframe (Wood et al., 2012; Wood et al., 2013). Indeed, the half-life of R121919 is short (∼2-3 hours, dependent on route of administration; Gehlert et al., 2007), allowing for repeated dosing across several days. Selective receptor occupancy for CRFR1a in the brain has been confirmed using peripherally administered doses between 2.5 mg/kg to 20 mg/kg, 1-hour after R121919 treatment (Heinrichs et al., 2002). To test the potential for CRFR1a to induce enduring neuroplasticity, offspring were again evaluated on these behavioral metrics at >P90 one week prior to brain tissue collection by P100. Each measure included only one male and one female from each litter (**see Table 1**).

**Table 1.** Total number of animals and litters included on each measure.

| Treatment | Number of Offspring<sup>a</sup> Included in Adolescent Behavior Analyses | Number of Offspring<sup>a</sup> Included in Adult Behavior Analyses | Number of Offspring<sup>a</sup> Included in Adult RNASeq Analyses |
| --- | --- | --- | --- |
| Saline | Male: 8<br>Female: 8 | Male: 8<br>Female: 8 | Male: 3<br>Female: 3 |
| CRFR1a | Male: 8<br>Female: 8 | Male: 8<br>Female: 8 | Male: 3<br>Female: 3 |

CRFR1a: corticotropin-releasing factor receptor 1 antagonist
<sup>a</sup>Only one male and one female offspring were evaluated from each litter.

### Offspring Behavioral Analyses

#### Locomotor Behavior

Prior to the social preference test (described below) animals were habituated to the test arena (40 cm x 40 cm x 28 cm) and video recorded for 5 minutes. To ensure that general ambulatory functioning was not impaired due to the CRFR1a treatment, locomotor activity (on P30 & >P90) was evaluated from the recording using automated behavioral monitoring software (Any-maze, Wood Dale, IL) to record distance travelled (m; Yan & Kentner, 2017; Connors et al., 2014; Núñez Estevez et al., 2020). Given that direct stimulation of CRFR1 in the extended amygdala can drive locomotor behavior (Chang et al., 2022), and early life alterations to amygdala CRFR expression is associated with later life mobility (Carty et al., 2010), we were also interested in whether early life CRFR1a exposure could result in either hyper- or hypo-activity in adulthood.

#### Social Preference Test

Sustained changes in adult social behavior have been previously associated with pubertal exposure to CRFR1a under non-stressful conditions (Veenit et al., 2014). To determine if we could replicate this effect, immediately following evaluation of locomotor activity, a five-minute social preference test (P30 & >P90) was videorecorded and later scored by trained observers (ODLog™ 2.0, http://www.macropodsoftware.com/) blinded to the group designations. The social preference test consisted of two cleaned wire containment cups placed on each end of the arena. One cup held a novel untreated rat of the same sex, age, size, and strain and the other cup held a novel object. The location of novel rats (n = 10) and novel objects was interchanged between trials. Rats were scored as actively investigating when their nose was directed within 2 cm of a containment cup, or it was touching the cup. A social preference index was calculated by the equation ([time spent with the rat] / [time spent with the inanimate object + time spent with the rat]) − 0.5; Scarborough et al., 2020) and the number of visits to the novel rat was recorded.

#### Social Discrimination Test

Twenty-four hours later (P31 & >P91), animals were placed back into the arena for an evaluation of social discrimination. This test was run and scored in an identical manner to the social preference test except both cups held an untreated rat of the same sex, age, size, and strain. In one cup was a novel rat, while the second cup held the familiar rat the experimental animal had been introduced to the day before. A social discrimination index was similarly calculated by the equation ([time spent with the novel rat] / [time spent with the familiar + time spent with the novel rat]) − 0.5).

#### Prepulse inhibition (PPI)

CRF and CRFR1a functioning in the amygdala are associated with PPI (Liang et al., 1992; Sutherland et al., 2010; Rajbhandari et al., 2015; Bijlsma et al., 2011). For this test, animals were placed into acoustic startle chambers (San Diego Instruments, San Diego, CA, USA) and allowed to habituate for 300 sec prior to the test session. A 65 dB background noise was present throughout the habituation period and test session, even when there was no presentation of a stimulus during a trial. During stimulus trials, rats were exposed to a 40 ms pulse of 120 dB white noise with or without the presentation of a prepulse. Prepulse intensity consisted of one of three intensity types: 8, 12, or 16 dB greater than the background noise. No-stimulus, pulse-alone, and prepulse-plus-pulse trials were each pseudorandomly presented 10 times, and the average trial interval was 15 ± 5 seconds. The % PPI for each prepulse intensity was calculated using the following formula: 1-(mean reactivity on prepulse-plus-pulse trials / mean reactivity on pulse alone trials) × 1/100 (Giovanoli et al., 2013).

#### Von Frey Test

CRFR signaling in the amygdala is associated with nociceptive responses in rats (Cui et al., 2004; Tian et al., 2024). To test whether early life exposure to the CRFR1a affected mechanical allodynia, animals were habituated inside an acrylic cage with a wire grid floor for 30-minutes. The von Frey test was evaluated using a pressure-meter which consisted of a hand-held force transducer fitted with a 0.7mm^2^ polypropylene tip (electronic von Frey anesthesiometer, IITC, Inc, Life Science Instruments, Woodland Hills, CA, USA). The tip was applied to the animal’s left hind paw with an increasing pressure until a flexion reflex occurred. The threshold of the applied weight (grams) was automatically recorded by the electronic pressure-meter when the animal withdrew its paw. The average of four test trials was calculated as the mechanical allodynia threshold (Yan & Kentner, 2017).

### Tissue Collection

On approximately P100, a week after the adult behavioral tests, offspring were anesthetized with a mixture of ketamine/xylazine (150 mg/kg /50 mg/kg, i.p.) between 1pm and 4pm. After intracardial perfusion with PBS, brains were removed, placed over ice and whole amygdala dissected using a brain block (Jia et al., 2012). Samples were frozen on dry ice and stored at −80°C until further processing.

### RNA Sequencing

Adult amygdala tissue samples were homogenized (Bead Ruptor 12, Omni International, Kennesaw, GA, USA) and the RNAeasy Plus Micro kit (Qiagen, Valencia, CA, USA) was used to extract RNA. Concentrations of RNA were evaluated with an Invitrogen Qubit 4 Fluorometer (Thermo Fisher Scientific, Inc, Waltham, USA) and the RNA Nano 6000 Assay Kit of the Agilent Bioanalyzer 2100 system (Agilent Technologies, CA, USA) was used to measure RNA integrity (RIN >7.9 = excellent quality). Following RNA-isolation (see Martz et al., 2024), amygdala samples (n = 3 per group) were more broadly analyzed using RNA Sequencing (RNA-seq) by Azenta Life Sciences (South Plainfield, NJ, USA) as described previously (Martz et al., 2024). An Agilent TapeStation (Agilent Technologies, Palo Alto, CA, USA) system was used to run the library and an Illumina NovaSeq 6000 (Illumina, San Diego, CA, USA) was used to sequence the samples according to the manufacturer’s instructions using a 2x150bp Paired End (PE) configuration (see Martz et al., 2024). The CLC Genomics Workbench v.23.0.2 software (QIAGEN Digital Insights, Aarhus, Denmark) was used for differential expression (DE) analysis and genes with an absolute fold change (FC) > 1.5, a Benjamini–Hochberg corrected p-value < 0.05, and false discovery (FDR) value < 0.05, were considered significantly DE. Volcano plots were constructed using the log_2_fold change and -log_10_ p-value in GraphPad Prism v.9.5.1. Heatmaps were generated using the free browser program Morpheus (https://software.broadinstitute.org/morpheus). Rank-rank hypergeometric overlap (RRHO) was used as a threshold-free approach to determine the extent of concordance/discordance between transcriptomic datasets (Cahill et al., 2018).

### Statistical Analyses

Statistical Software for the Social Sciences (SPSS) was used to perform statistical analyses. To keep in line with the National Institutes of Health (NIH) policy to consider sex as a biological variable (Clayton et al., 2018), we include both males and females in our study but did not set out to evaluate sex differences. Therefore, since sex was not compared, t-tests were applied to the behavioral data as appropriate, except for the PPI and von Frey tests. %PPI, like body weight, was evaluated using repeated measures ANOVA. In the former case, repeated measures ANOVA was performed to compare differences in %PPI across each trial type. Baseline startle reactivity was skewed; therefore, a ln-transformation was applied prior to analysis (Csomor et al., 2008). Body weight was used as a covariate for the evaluation of mechanical allodynia in the von Frey test using ANCOVA (Yan & Kentner, 2017). The partial eta-squared (*n_p_*^2^) is reported as an index of effect size (Miles & Shevlin, 2001). All data are expressed as mean ± SEM.

RNA sequencing data were analyzed as previously described (Martz et al., 2024). Briefly, DESeq2 was employed to identify differentially expressed (DE) genes based on a p < 0.05, Benjamini–Hochberg false discovery rate corrected (FDR) and fold change (FC) > 1.5 and RRHO2 was used as a threshold-free approach to determine the extent of concordance/discordance between transcriptomic datasets (Cahill et al., 2018). Heatmaps were generated using the MultiExperiment Viewer (National Library of Medicine, USA) and gene ontology was determined using the Database for Annotation, Visualization and Integrated Discovery functional annotation cluster tool (https://david.ncifcrf.gov/).

## Results

### CRFR1 antagonism has acute effects on adolescent behavior

Repeated subchronic administration of CRFR1a, R121919 did not affect body weight in either male or female adolescent rats (p>0.05, **Supplemental Fig. 1A,B**). Similarly, acute CRFR1 antagonism did not impact adolescent locomotor activity, social preference, the number of visits made to a social conspecific (p>0.05, **Fig. 2A,B,C**), social discrimination, mechanical allodynia, nor was baseline startle reactivity affected (p>0.05, **Supplemental Fig. 2A,B, C**). Systemic blockade of CRFR1 did however interrupt %PPI, which was used to assess sensorimotor gating. A significant main effect of CRFR1a showed attenuated %PPI response across the 77 dB and 81 dB prepulse intensities in male adolescents (77 dB: F(1, 14) = 10.174, p = 0.007, n_p_^2^ = 0.439; 81 dB: F(1, 14) = 8.080, p = 0.014, n_p_^2^ = 0.383, **Fig. 2D**). Mean %PPI was calculated by collapsing %PPI across all dB intensities. There was a main effect of CRFR1 antagonism, where mean %PPI was reduced in males exposed to R121919 (t(14) = 2.822, p = 0.007, n_p_^2^ = 0.363, **Fig. 2E**).

**Figure 2.**
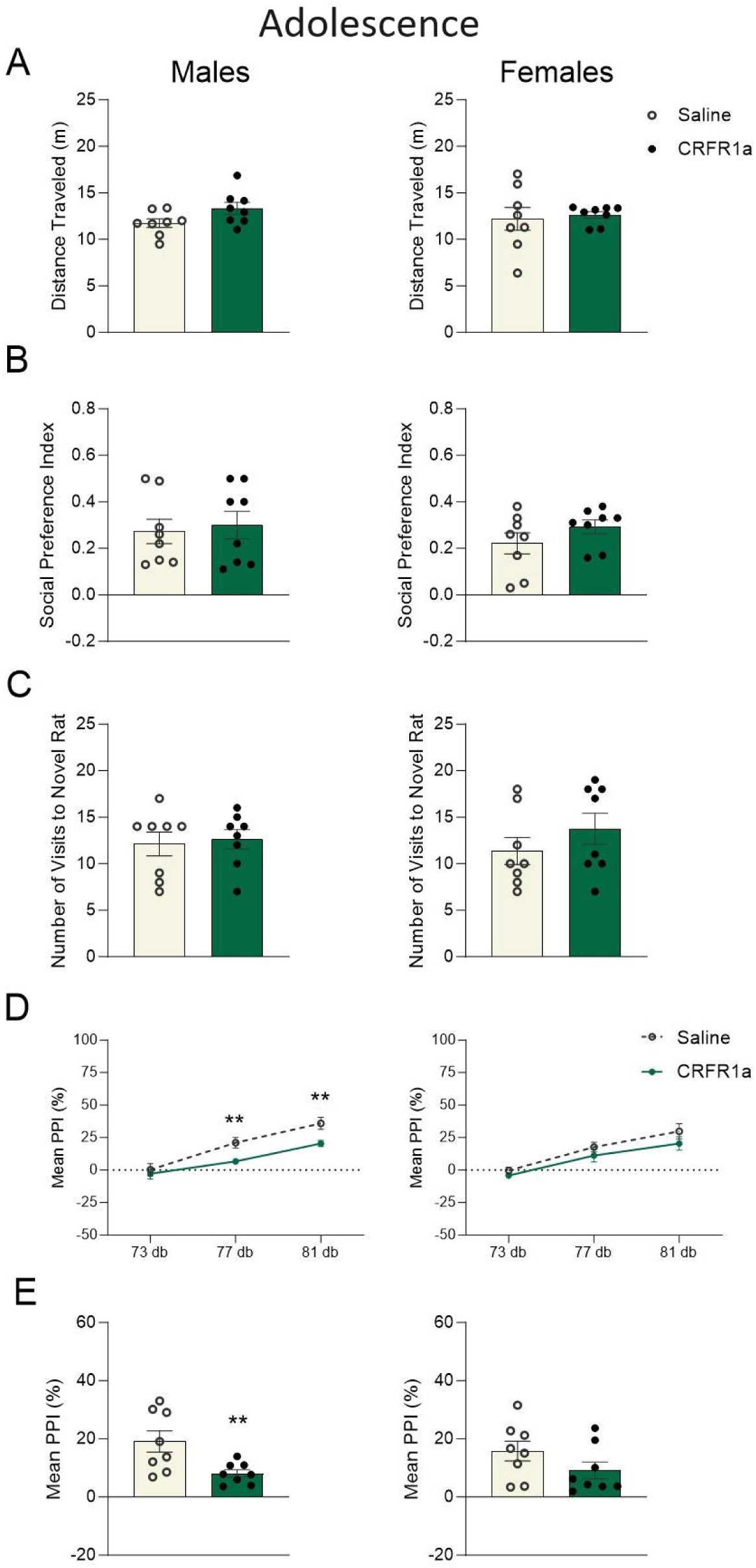
Adolescent behavioral functioning is altered in a sex-specific manner following acute CRFR1 antagonism. Graphs display male (left) and female (right) data for adolescent A) total distance traveled (m), B) social preference index, C) total number of visits to a novel rat in the social preference test, D) percent prepulse inhibition (PPI%) across each trial type, and (E) mean %PPI collapsed across all trial types. Data are expressed as mean ±SEM. *p < 0.05, **p < 0.01, Saline versus adolescent CRFR1 antagonism, n = 8.

### CRFR1 antagonism during peripuberty has sustained effects on behavioral plasticity

Acute peripubertal CRFR1a exposure was associated with increased locomotor activity in female adult rats (t(14) =-3.267, p = 0.006, n_p_^2^ = 0.443, **Fig. 3A**). While the social preference index was not affected (p>0.05, **Fig. 3B**), adult females exposed to pubertal CRFR1a displayed a higher number of visits to their novel conspecifics (t(14) = -2.171, p = 0.048, n_p_^2^ = 0.252, **Fig. 3C**). This effect appeared to be specific to the social component of the test as female animals did not differ in the number of visits to the novel object (p>0.05; Female Saline: 4.13±0.40 versus Female CRHR1a: 4.50±0.73; Male Saline: 7.38±1.10 versus Male CRFR1a: 6.00±0.60). Moreover, adolescent CRFR1a was associated with a modest, yet persistent reduction in %PPI in adult male and female rats, despite there being no active drug exposure at the time of testing (males (81 dB): F(1, 14) = 5.510, p = 0.034, n ^2^ = 0.282; females (73 dB): F(1, 14) = 7.028, p = 0.019, n_p_^2^ = 0.334, **Fig. 3D**). Mean %PPI (p > 0.05, **Fig. 3E**), baseline startle reactivity, social discrimination, and mechanical allodynia were not affected in adulthood for either male or female rats (p > 0.05, **Supplemental Fig. 2D, E, F**).

**Figure 3.**
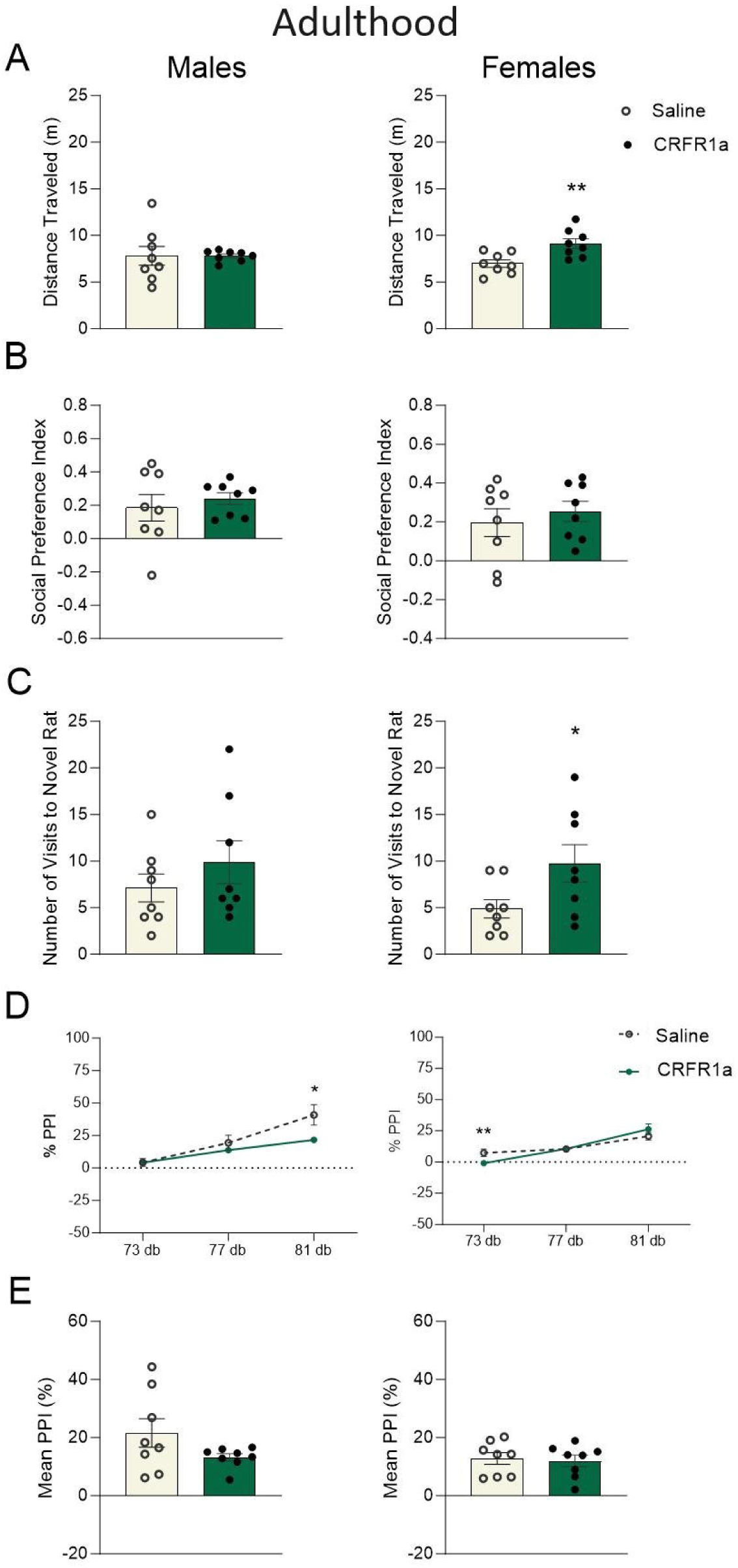
Adolescent exposure to CRFR1 antagonism induced prolonged and sex- specific effects on behavioral plasticity. Graphs display male (left) and female (right) data for adult A) total distance traveled (m), B) social preference index, C) total number of visits to a novel rat in the social preference test, D) percent prepulse inhibition (PPI%) across each trial type, and (E) mean %PPI collapsed across all trial types. Data are expressed as mean ±SEM. *p < 0.05, **p < 0.01, Saline versus adolescent CRFR1 antagonism, n = 8.

### Peripubertal CRFR1 antagonism imparted lasting effects on adult neurophysiology and neuroplasticity markers

RNA sequencing was performed on whole amygdala tissue from adult animals to evaluate the long-term transcriptional effects of peripubertal CRFR1a exposure. In males, 219 genes were significantly DE by CRFR1a (q<0.05 versus Saline; **Fig. 4A**) and 31 genes were significantly DE by CRFR1a exposure in females (q<0.05; **Fig. 4B**). Threshold-free RRHO analysis showed substantial overlap in gene expression profiles across males and females as a function of CRFR1a exposure (**Fig. 4C**). **Fig. 5** shows the top pathways affected by peripubertal CRHR1a and include pathways related to behavior, such as learning and memory, stress responses, limbic system development, and regulation of hormone levels in males and central nervous system myelination, cell junction organization, and glutamatergic regulation of synaptic transmission in females. **Supplemental Results Tables 1, 2, and 3** show specific gene expression changes in each sex. Notably, the modulation of chemical synaptic transmission was affected by CRHR1a in both sexes. Heatmap clustering identified alterations in genes belonging to long-term potentiation (LTP; **Fig. 4D**) and long-term depression (LTD; **Fig. 4E**), in addition to stress (**Fig. 4F**), dopamine (**Fig. 4G**), and serotonin (**Fig. 4H**) pathways in peripubertal CRFR1a exposed males and females. Pathways related to glucocorticoid binding, oxytocin, cannabinoids, glutamate/GABA, parvalbumin, and microglia were also identified (**Supplemental Fig. 3A-F**). For more information on heatmap clustering, see **Supplemental Results Table 1**. Threshold-free RRHO analysis showed a high degree of overlap between the effect of sex in controls and the effect of sex in CRFR1a exposed rats (**Supplemental Fig. 4**).

**Figure 4.**
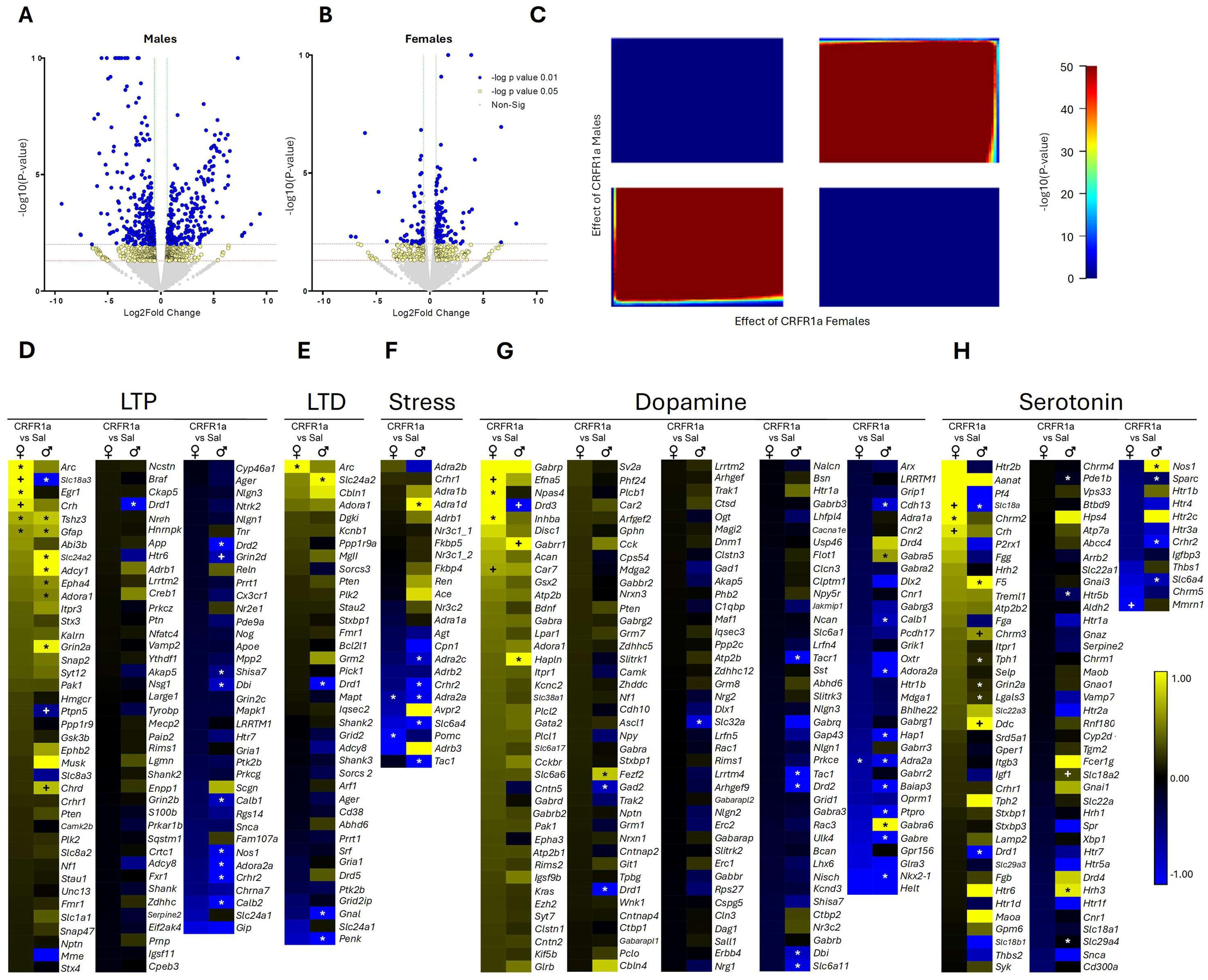
Transcriptomic analyses of amygdala tissue samples collected from adult male and female animals following peripubertal exposure to CRFR1a. Volcano plots for differentially expressed (DE) transcripts in A) male (1064 genes) and B) female (31 genes) rats, with -log_10_ pvalue on the y-axis and log_2_ FC on the x-axis. Each dot represents a gene, with colors indicating the magnitude of significance (yellow = p < 0.05, FC > 1.5; blue = p < 0.01, FC > 1.5). Threshold-free RRHO analysis indicates that the transcriptional alterations that occur as a function of C) peripubertal CRFR1a exposure largely overlap between males and females. Heatmaps of D) long-term potentiation (LTP), E) long-term depression (LTD), F) stress, G) dopamine, and H) serotonin related genes, with effects represented as FC between Saline and CRFR1a exposed males, and between Saline and CRFR1a exposed females. *Genes that reached the Benjamini-Hochberg corrected effect of 20%. +Genes with p < 0.05 and FC > 1.5.

**Figure 5.**
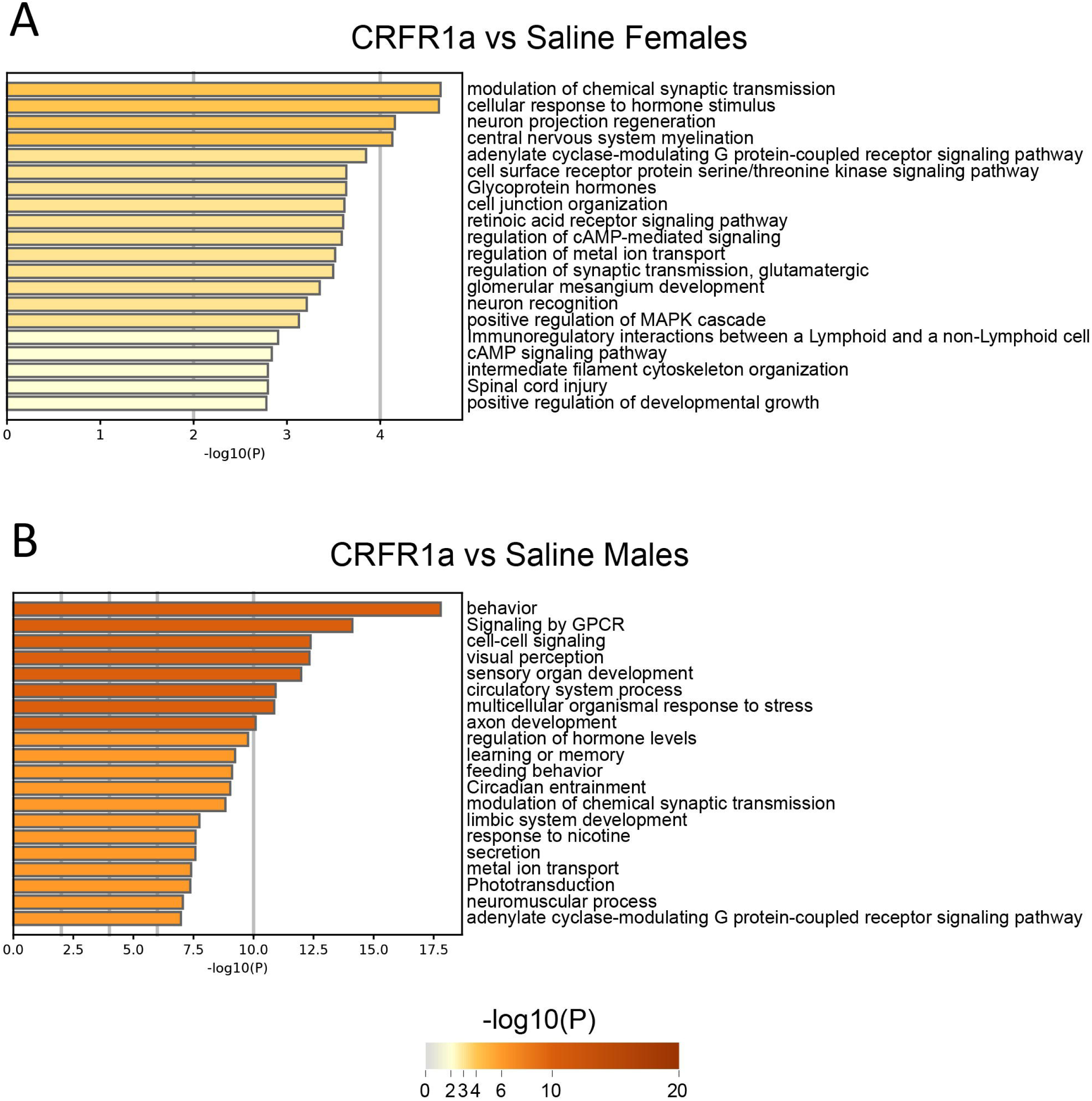
Pathways of top differentially expressed transcripts after peripubertal CRFR1 exposure. A) Top pathways represented by transcripts differentially expressed in females (A) and males (B) as a function of CRFR1a.

## Discussion

In the present study, we replicate and extend on evidence demonstrating that temporary blockade of CRFR1 during the peripubertal period induces long-term programming effects on adult behavior (Veenit et al., 2014). Our own findings suggest that this behavioral programming can occur in both male and female rats and is likely due to persistent changes in gene pathways associated with neural plasticity and stress. Injections of the CRFR1a R121919 immediately reduced %PPI in adolescent males, but not females. This suggests that adolescent male rats may be more sensitive to the immediate effects of this drug. In adulthood, CRFR1a exposed males continued to show deficits in PPI, while alterations in locomotion, PPI, and social behavior emerged in CRFR1a females, in comparison to saline controls. Although the behavioral programming effects of CRFR1 antagonism were more evident in the adult female dataset, transcriptomic data revealed that males experienced many gene expression changes in the amygdala. Overall, these findings identify peripuberty as a sensitive period for disruption of CRFR1 signaling, even in the absence of a stressor.

Although this work did not investigate the effects of CRFR1 blockade in the context of stress, adolescent administration of R121919 had long-term effects on the expression of genes in the amygdala related to stress and anxiety. CRFR2, encoded by the gene *Crhr2*, performs functions distinct from those facilitated by CRFR1, but the two receptors work in concert to orchestrate stress responses (Kishimoto et al., 2020; Vasconcelos et al., 2020). In our males, temporary blockade of CRFR1 during the peripubertal period caused long-term downregulation of *Crhr2* in adulthood, indicating that stress signaling in the amygdala was permanently affected. Indeed, pathways related to multicellular organismal responses to stress and the regulation of hormone levels were both altered in our CRFR1a adult males. In addition to its role in stress signaling, the amygdala is an important modulator of behaviors involving social tasks and sensorimotor gating (Ko, 2017; Wan and Swerdlow, 1997; Cano et al., 2021), which were disrupted in our pubertal exposed CRFR1a animals when they reached adulthood. These sustained alterations in behavior demonstrate behavioral plasticity following the early life CRFR1a challenge. Notably, CRFR1 is densely expressed in the amygdala (Dedic et al., 2018; Wolfe et al., 2019) and has been implicated in the modulation of social behavior and PPI in rodents (Hostetler and Ryabinin, 2013; Risbrough et al., 2004; Groenink et al., 2008; Sutherland and Conti, 2011). In a social preference task, adult female mice treated with CRFR1a 1 hour before testing spent less time with a novel mouse compared to a novel object. In contrast, CRFR1a males had a stronger preference for their novel conspecific (Piccin and Contarino, 2020). In our study, CRFR1a was administered to peripubertal rats 1 hour before a social preference task and had no immediate effects. It is unclear if the discrepancy in our findings compared to those of Piccin and Contarino (2020) is due to the difference in species, CRFR1a drug, the disparate ages of the animals, or a combination of these factors. When our animals were tested in adulthood, although there was still no effect on social preference, adult CRFR1a females surprisingly made a greater number of visits to the novel rat than did saline controls. Data from the amygdala revealed that pathways with genes previously implicated in sociability (Harris et al., 2016; Froemke and Young, 2021; Ike et al., 2023) were altered in our adult CRFR1a males, but not CRFR1a females. This suggests that male rats may have experienced some neural compensation, protecting them against CRFR1a-induced changes in adult social behavior. However, our males, similar to the females, were vulnerable to long-term changes in %PPI. Taken together, sex differences in amygdalar function may underlie the discrepancies in social behavior outcomes following adolescent CRFR1a exposure and future work will need to test this directly.

With respect to sensorimotor gating, 1) adult male rats that received infusions of CRF or 2) adult male mice that were bred to overexpress CRF each exhibited reductions in PPI that were rescued by CRFR1 antagonism (Risbrough et al., 2004; Groenink et al., 2008). Conversely, Sutherland and Conti (2011) found that PPI was not altered by CRFR1a in adult male rats that had experienced restraint stress, suggesting that the effects of CRFR1a on PPI may vary based on stress experience and physiological state. Here we show that peripubertal administration of a CRFR1a, under basal conditions, reduced PPI both acutely and chronically for male rats. Analysis of the female data showed that they were only affected in adulthood. Divergent sensorimotor gating behaviors have been associated with neurodevelopmental disorders such as autism and schizophrenia (Perry et al., 2007; San-Martin et al., 2020), and we found that pubertal exposure to CRFR1a altered gene pathways in the amygdala related to these disorders; Herrero et al., 2020; Caubit et al., 2016; Sohal and Rubenstein, 2019; Boiko et al., 2020). Among the males that received the CRFR1a, expression of four genes related to GABAergic signaling (*Gabra6, Gabra5, Gabre, and Gad2*) were dysregulated, while expression of one GABA-related gene (*Gabrg1*) was altered among CRFR1a females. GABAergic neurons act in the amygdala to inhibit inappropriate behavioral responses, so disruptions in this system could lead to detrimental changes in behavior (Jie et al., 2018), particularly with respect to the sensorimotor responses observed in the present study. In sum, dysregulation of gene pathways combined with the sustained changes in PPI in our CRFR1a-treated rats are indicative of permanent changes to stress systems in the brain

Relatedly, changes in neural plasticity induced by CRFR1a have previously been shown, but in the context of subsequent exposure to stress (Ivy et al., 2010; Short et al., 2020). Here, we find that gene pathways influenced by CRFR1 exposure relate to synaptic plasticity and metaplasticity, which, as a family, underwent the most pronounced changes in expression. Although we did not assess fear behavior or learning and memory, it is likely that peripubertal blockade of CRFR1 would influence adult fear and memory formation/consolidation, in addition to the alterations to PPI and social behavior observed here, based on the identified gene pathways altered in the amygdala of our CRFR1a animals (Fig. 5).

With respect to the data interpretation of the current study, it is important to consider some methodological limitations. First, in line with the 3Rs of animal research’s principle of Reduction (NC3Rs, n.d), our animals were siblings of offspring from gestationally treated saline control dams utilized in a previous study (Martz et al., 2024). Therefore, it is possible that our effects may not replicate in offspring from untreated dams. Indeed, intraperitoneal injections can be stressful (although minimally; Al Shoyaib et al., 2019) and the CRF system can be affected by prenatal stress experiences (Cratty et al., 1995; Mueller & Bale, 2008; Zhao et al., 2021). Additionally, the susceptibility of offspring to demonstrate changes in brain and behavior because of pubertal CRFR1 antagonism could be due to the use of a single time-period of drug exposure e.g., (P30-P33) or to differences in the timing of pubertal transitions across male and female rodents (Ghasemi et al., 2021). Because our animals underwent significant behavioral testing, it must also be considered whether the influence of CRFR1a on gene expression was a function of their experience across our study groups. That said, we did include saline vehicle counterparts to our male and female CRFR1a groups, to control for experience. It is unclear how translationally informative gene expression changes would be from behaviorally naive animals with limited environmental experiences (see Kentner et al., 2021). Additionally, for our PPI protocol, only a single high intensity stimulus (120 dB) was used. Although this is typical in many PPI studies, this could limit the interpretation of our findings.

In summary, this work identifies peripuberty as a malleable period of development, during which time the brain is sensitive to discrete and temporary changes in CRF signaling activity, leading to sustained behavioral changes, even in the absence of a stressor. Blockade of CRFR1 across 4 days during early adolescence had long-lasting effects on synaptic plasticity and other neural domains, including dysregulation of genes implicated in neurodevelopmental disorders and stress. Notably, these dysregulated genes directly relate to the long-term alterations in PPI and social behavior reported here. While many studies have shown that too much stress during adolescence negatively impacts neurodevelopment (Carr et al., 2013), it is also known that too little stress can have negative effects (Kirby et al., 2013). Here we see that an acute influence on central stress systems during early adolescence has long-lasting effects on neural and behavioral outcomes. These findings have important implications for the use of psychoactive treatments during the peripubertal period, given their potential for far-reaching and sustained impacts on brain development and behavior.

---

**Supplementary Figure 1. Adolescent and adult body weight data following CRFR1 antagonism.** Graphs display male (left) and female (right) data for adult A) adolescent and B) adult body weights (g). Data are expressed as mean ±SEM. *p < 0.05, **p < 0.01, Saline versus adolescent CRFR1 antagonism, n = 8.

**Supplementary Figure 2. Adolescent and adult behavioral measures following CRFR1 antagonism.** Graphs display male (left) and female (right) data for adolescent (top panel) and adult (bottom panel) A,D) social discrimination, B,E) baseline startle reactivity [ln-transformed] (mV) on the PPI test, and C,F) mechanical allodynia threshold (g) on the von Frey test. Data are expressed as mean ±SEM. *p < 0.05, **p < 0.01, Saline versus adolescent CRFR1 antagonism, n = 8.

**Supplemental Figure 3. Effect of peripubertal CRFR1a exposure on transcriptomic analyses of amygdala tissue samples collected from adult male and female offspring.** Heatmaps of A) glucocorticoid binding (GR)-, B) oxytocin-, C) cannabinoid (CB)-, D) glutamate (Glu)/GABA, E) parvalbumin (PV)-, and F) microglia-related genes, with effects represented as fold change (FC) between CRFR1a and Saline exposed males, and between CRFR1a and Saline exposed females. *Genes that reached the Benjamini-Hochberg corrected effect of 20%. +Genes with p<0.05 and FC>1.5.

**Supplemental Figure 4. Threshold-free RRHO analysis showed a high degree of overlap between the effect of sex in controls and the effect of sex in CRFR1a exposed rats.**

**Supplemental Results Table 1. Gene Expression in Saline vs. CRFR1a Amygdala**

**Supplemental Results Table 2. Females CRFR1a vs Sal**

**Supplemental Results Table 3. Males CRFR1a vs Sal**

## Author Contributions

J.M., T.J.L., S.S., & M.A.S., ran the experiments; J.M., M.A.S., M.L.S., & A.C.K. analyzed and interpreted the data; J.M and A.C.K. wrote the manuscript; J.M., M.A.S., M.LS., T.J.L, S.S, & A.C.K., edited the manuscript; A.C.K., designed and supervised the study.

