## Supplementary figures and images for "Peripubertal antagonism of corticotropin-releasing factor receptor 1 results in sustained changes in behavioral plasticity and the transcriptomic profile of the amygdala"

### Supplemental Figure 1

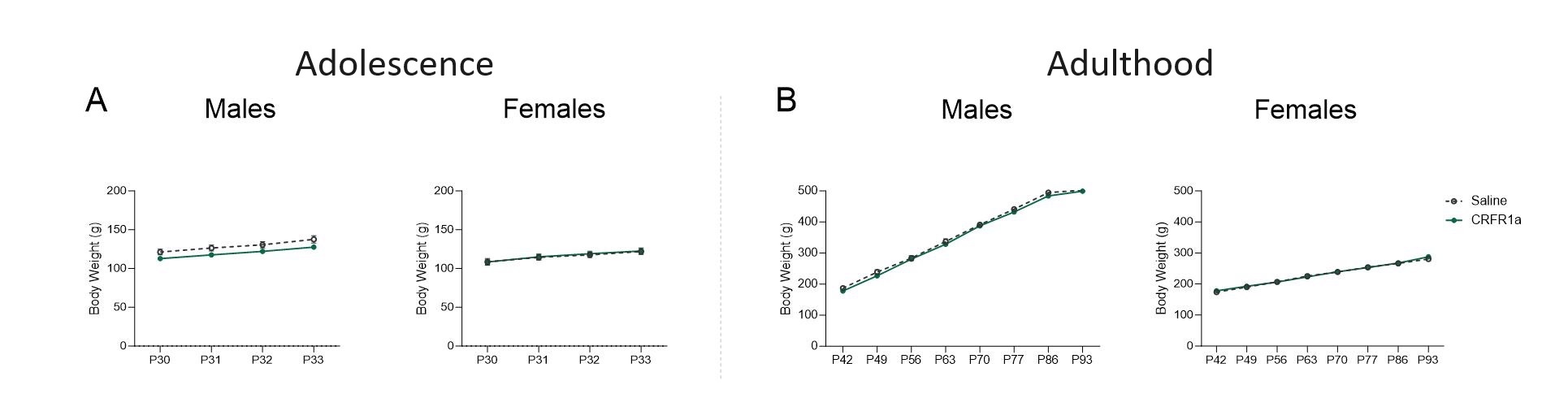

### Supplemental Figure 2

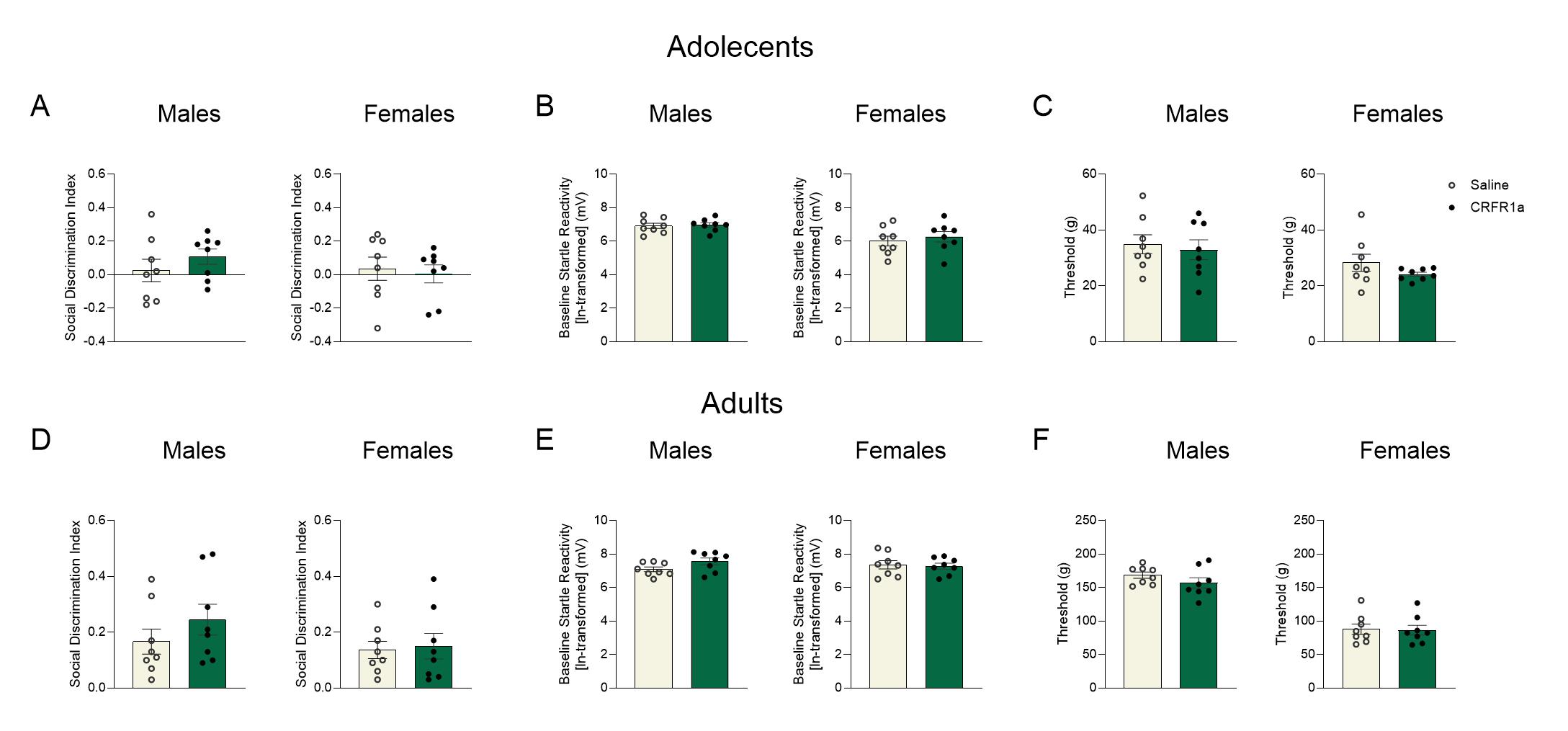

### Supplemental Figure 3

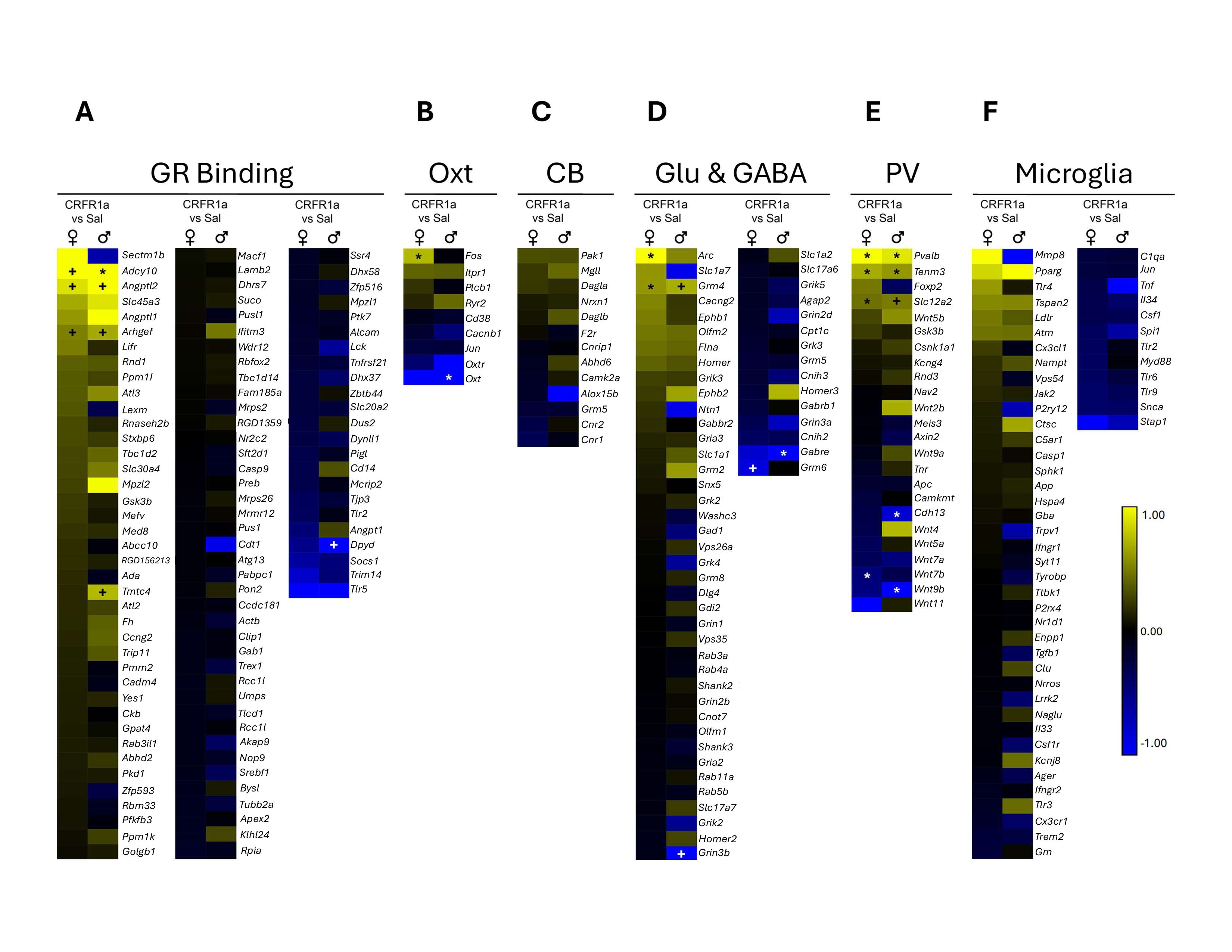

### Supplemental Figure 4

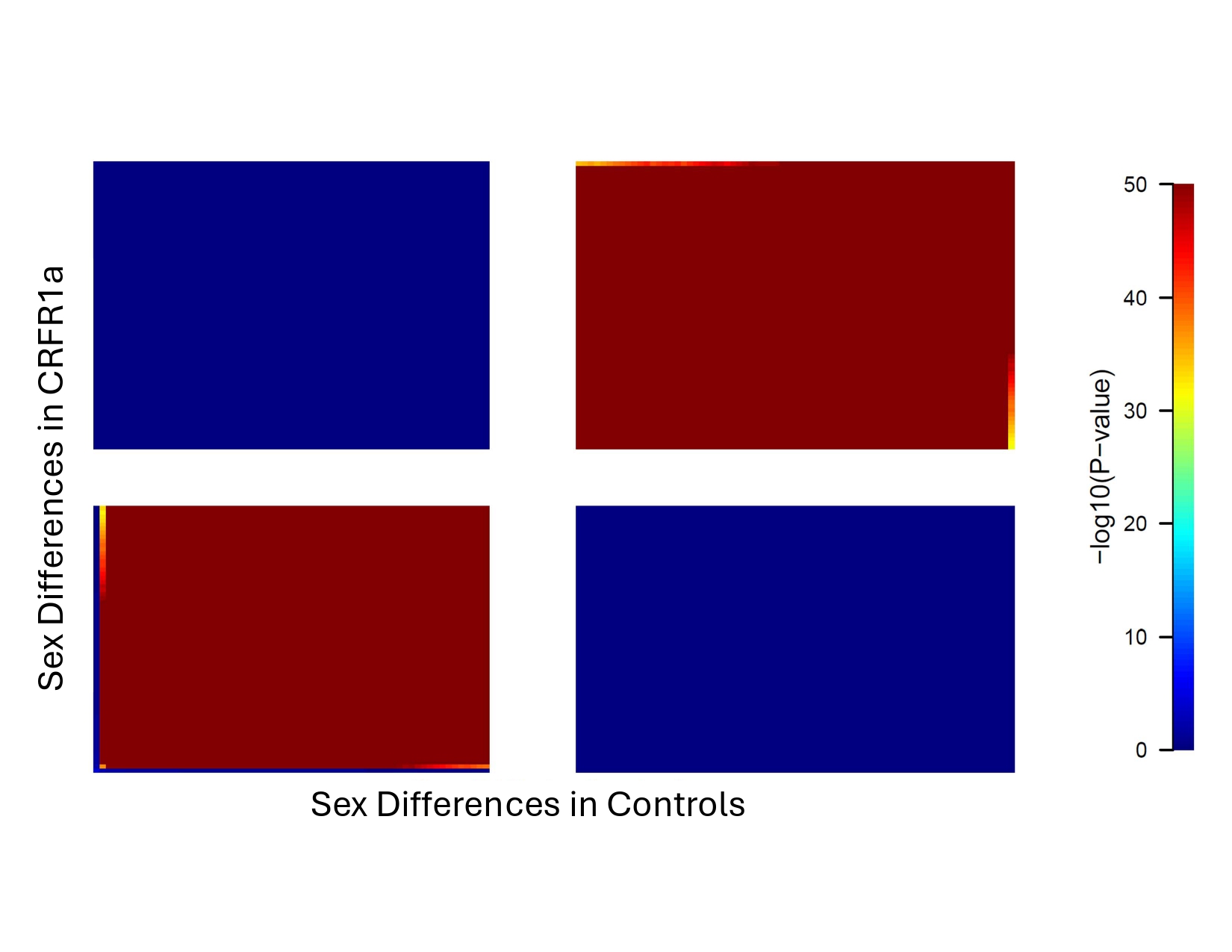
